# Detection and characterisation of alpha- and betacoronaviruses in rodents and bats from Germany, France, Belgium and Ireland

**DOI:** 10.64898/2026.09.23.753110

**Authors:** Elena Sgarabotto, Vinaya Venkat, Eda Altan, Daan Gyzels, Mert Erdin, Veronica Lehtinen, Thanakorn Niamsap, Sofia Greilich, Akseli Valta, Maxime Galan, Jasmin Firozpoor, Jana. A. Eccard, Jens Jacob, Christian Imholt, Vincent Sluydts, Valeria Colombo, Herwig Leirs, Andrew McManus, Peter Stuart, Sophie Gryseels, Luc De Bruyn, Christian C. Voigt, Uwe Hoffmeister, Hussein Alburkat, Vincent Bourret, Teemu Smura, Guillaume Castel, Nathalie Charbonnel, Tarja Sironen, Lara Dutra

## Abstract

Recent zoonotic coronavirus outbreaks have sparked an interest in understanding coronaviruses circulating in animal reservoirs. While bats are recognised as major reservoirs for these viruses, coronaviruses have also been detected in a wide range of terrestrial small mammals, particularly rodents. However, coronavirus diversity in these mammals, specifically in Europe, is still poorly characterised.

In this study, we detected and characterised coronaviruses in terrestrial small mammals and bats from Belgium, France, Germany, and Ireland. We screened tissue samples from rodents and shrews and environmental bat guano using pan-coronavirus real-time and conventional PCRs, with positive results confirmed by sanger sequencing. Among 2667 bat and 1152 terrestrial mammal samples, we detected 59 positive samples, 36 from bats and 23 from rodents, from which we recovered 60 coronavirus sequences, spanning both *Alphacoronavirus* and *Betacoronavirus* lineages. Through next generation sequencing we identified one *Pipistrellus pygmaeus* bat guano sample carrying two distinct alphacoronaviruses, representing Nyctacovirus and Pedacovirus subgenera.

Among rodents, most coronavirus-positive samples originated from a zoo in Belgium, where several CoV lineages co-circulated. The identified rodent coronaviruses clustered among previously described rodent-associated lineages, some of which also include human and other mammal-derived coronavirus sequences. Consistent with previous studies, we found sarbecoviruses in horseshoe bats (particularly *Rhinolophus ferrumequinum* and *R. hipposideros*) and merbecoviruses in *Plecotus auritus* bats, and these viruses are distantly related to the human coronaviruses of the respective families. Similar to bat coronaviruses, some rodent coronaviruses strains seemed to be association with specific rodent species.

In conclusion, our study enhances the understanding of coronaviruses in rodents and bats, revealing potential hotspots for these viruses and their implication for human and animal health. These findings underscore the necessity for continued surveillance of coronaviruses in wildlife reservoirs to mitigate future zoonotic spillover risks.

**Author Summary:** Following recent zoonotic outbreaks, there is growing interest in understanding wildlife reservoirs for coronaviruses. While bats are well-known carriers of these viruses, terrestrial small mammals, specifically rodents, have received less attention. As a result, our knowledge of the diversity of coronaviruses in rodent remains limited. Hence, we investigated the diversity of coronaviruses in bats from environmental guano and terrestrial small mammals from their tissues in Belgium, France, Germany and Ireland. We identified a diverse range of coronaviruses in both rodents and bats that related to other human and animal infecting coronaviruses, but this does not imply their immediate zoonotic potential. Additionally, we found that some bat and rodent coronaviruses seemed to be associated with particular host species. These finding broaden the current knowledge of the diversity of coronaviruses in European small mammals and highlight the role of including rodents alongside bats in wildlife surveillance to deepen our understanding of virus diversity and their associated potential zoonotic risks.

## Introduction

Long known for causing mild respiratory diseases in humans, and acute respiratory and gastrointestinal infections in livestock, coronaviruses (CoVs) have become pathogens of major concern in the last two decades. Although human CoVs (hCoVs), such as hCoV HKU1, NL63, OC43 and 229E, commonly circulate and cause mild seasonal respiratory infections, the spillover of three highly pathogenic viruses from wildlife to humans has sparked global interest in wildlife coronaviruses and their zoonotic potential^1^. The spillover events of SARS, MERS, and SARS-CoV-2, originating from bats and involving intermediate mammalian hosts, demonstrate how coronaviruses can cross species barriers to cause global outbreaks^2–5^.

Coronaviruses belong to the family *Coronaviridae,* subfamily *Orthocoronavirinae* and are divided into four genera: *Alphacoronavirus*, *Betacoronavirus*, *Gammacoronavirus* and *Deltacoronavirus*. While gamma-CoVs and delta-CoVs mostly infect birds, alpha-CoVs and beta-CoVs infect mammals. Bats have long been recognised as significant carriers of zoonotic pathogens such as Ebola virus, Hendra viruses, Nipah virus, Rabies virus and coronaviruses^6^ . The coronaviruses they carry also include ancestors of coronaviruses responsible for major human outbreaks^7^. Their unique immune system, high species diversity, large and densely structured colonies, wide ecological distribution, ability to fly and cross-colony interactions make them well-suited to sustain and transmit viruses^1,4,8–11^.

Alongside bats, other mammals such as rodents and shrews are known to host and spread coronaviruses, like the Mouse Hepatitis Virus (murine coronavirus)^12–16^. The hCoVs HKU1 and OC43 are related to the murine coronavirus and have likely originated from rodents^2,17^. Recent studies from various regions have uncovered additional more alpha- and beta-CoVs in terrestrial small mammals^18,19^. The social behaviour and widespread presence of these animals may facilitate the intraspecies and/or interspecies or zoonotic transmission of infectious agents through close contact or insect vectors^7^. Altogether, these studies suggest that rodents harbour a high diversity of coronaviruses and may serve as potential source of future virus spillovers^5,13–15,20–24^.

In Europe, coronaviruses have been detected in multiple bat species, and novel coronaviruses have been reported in small mammals (voles, squirrels, hedgehogs, shrews) in France, United Kingdom, Sweden, Germany, Portugal and Poland ^5,13–15,20–26^. However, the knowledge on overall diversity of coronaviruses in terrestrial small mammals - particularly rodents - remains limited. To address this gap, we investigate coronavirus presence in small mammals of Belgium, Germany, France and Ireland, to better characterise the coronaviruses they carry, identify the host species associated with the detected viruses and provide additional data on their occurrence in the sampled locations.

## Results

### Sample overview

To investigate the presence and diversity of coronaviruses among small mammals in Europe, we analysed 1152 intestinal tissues (colon) of terrestrial small mammals (Table 1) and 2667 bat guano samples (Table 2), from four European countries, Belgium, France, Ireland and Germany (Figure 1). The 1152 colon samples were mostly from rodents of the families Cricetidae and Muridae, along with a few red squirrels, *Sciurus vulgaris* (Rodentia: Sciuridae), and *Crocidura russula* (Eulipotyphla: Soricidae). The 2667 guano samples were collected from the environment and expected to be from bats of *Vespertilionidae*, *Rhinolophidae* and *Miniopteridae* families based on prior local bat monitoring studies (Table 2)^27–32^. Only CoV positive guano samples were barcoded.

**Table 1.** Distribution of rodent and shrew samples analysed for the presence of coronaviruses according to species and country of origin.

| Scientific name (Order: Family) | Common name | Belgium<br>(6 sites) | France<br>(6 sites) | Germany<br>(7 sites) | Ireland<br>(6 sites) | Total |
| --- | --- | --- | --- | --- | --- | --- |
| *Apodemus agrarius* (Rodentia: Muridae) | Striped field mouse | 0 | 0 | 15 | 0 | 15 |
| *Apodemus flavicollis* (Rodentia: Muridae) | Yellow-necked mouse | 0 | 141 | 63 | 0 | 204 |
| *Apodemus sylvaticus* (Rodentia: Muridae) | Field mouse | 130 | 120 | 0 | 122 | 372 |
| *Clethrionomys glareolus* (Rodentia: Cricetidae) | Bank vole | 146 | 151 | 79 | 99 | 475 |
| *Microtus arvalis* (Rodentia: Cricetidae) | Common vole | 0 | 2 | 1 | 0 | 3 |
| *Microtus subterraneus* (Rodentia: Cricetidae) | European pine vole | 0 | 1 | 0 | 0 | 1 |
| *Mus musculus* (Rodentia: Muridae) | House mouse | 5 | 18 | 0 | 0 | 23 |
| *Rattus norvegicus* (Rodentia: Muridae) | Brown rat | 19 | 15 | 0 | 0 | 34 |
| *Sciurus vulgaris* (Rodentia: Sciuridae) | Red squirrel | 2 | 0 | 0 | 0 | 2 |
| *Crocidura russula* (Eulipotyphla: Soricidae) | Greater white-toothed shrew | 10 | 13 | 0 | 0 | 23 |
| **Total** |  | 312 | 461 | 158 | 221 | 1152 |

**Table 2.**
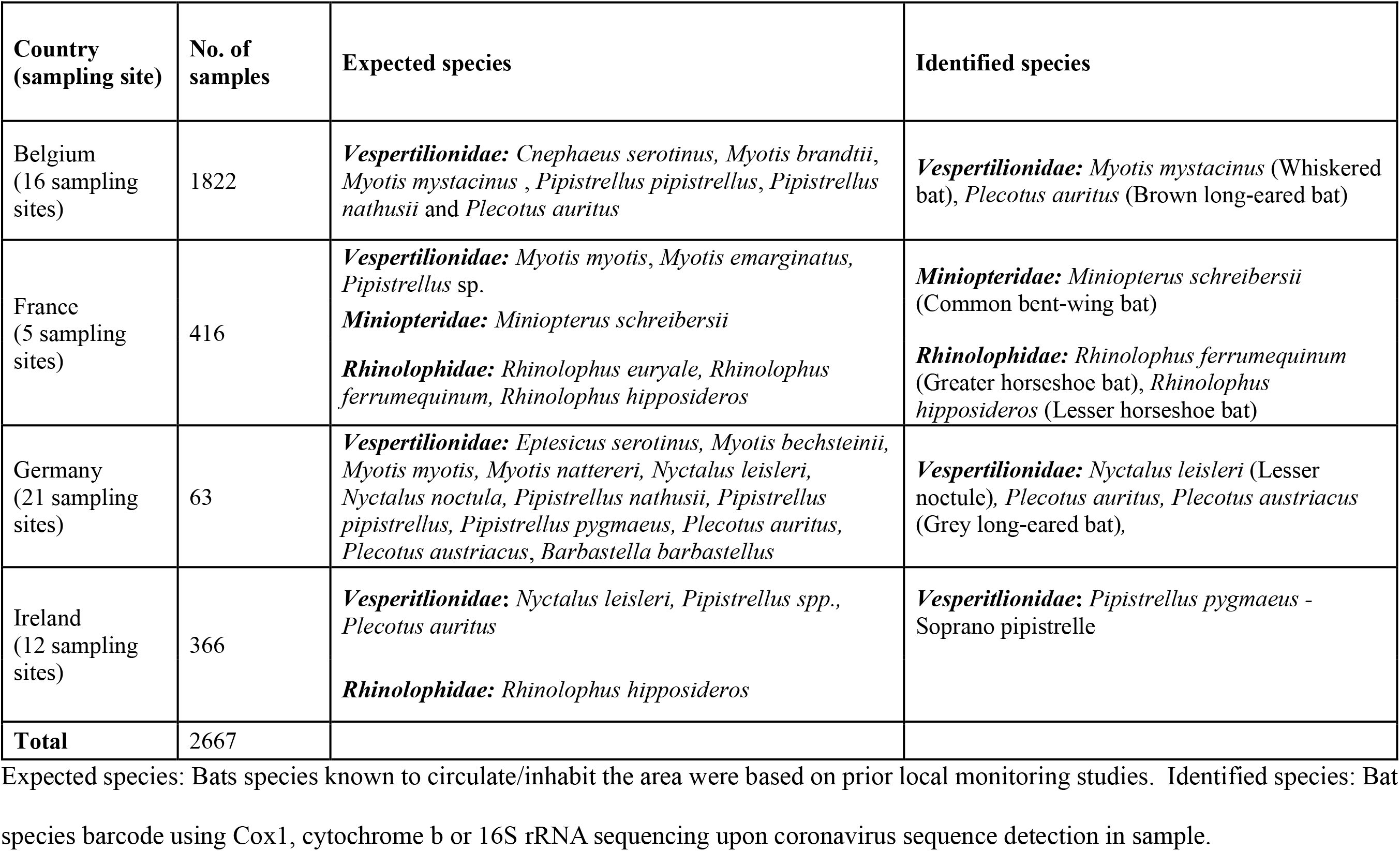
Distribution of guano samples analysed for the presence of coronaviruses according to country of origin and expected species.

| Country (sampling site) | No. of samples | Expected species | Identified species |
| --- | --- | --- | --- |
| Belgium (16 sampling sites) | 1822 | *Vespertilionidae*: *Cnephaeus serotinus*, *Myotis brandtii*, *Myotis mystacinus*, *Pipistrellus pipistrellus*, *Pipistrellus nathusii* and *Plecotus auritus* | *Vespertilionidae*: *Myotis mystacinus* (Whiskered bat), *Plecotus auritus* (Brown long-eared bat) |
| France (5 sampling sites) | 416 | *Vespertilionidae*: *Myotis myotis*, *Myotis emarginatus*, *Pipistrellus* sp.<br>*Miniopteridae*: *Miniopterus schreibersii*<br>*Rhinolophidae*: *Rhinolophus euryale*, *Rhinolophus ferrumequinum*, *Rhinolophus hipposideros* | *Miniopteridae*: *Miniopterus schreibersii* (Common bent-wing bat)<br>*Rhinolophidae*: *Rhinolophus ferrumequinum* (Greater horseshoe bat), *Rhinolophus hipposideros* (Lesser horseshoe bat) |
| Germany (21 sampling sites) | 63 | *Vespertilionidae*: *Eptesicus serotinus*, *Myotis bechsteinii*, *Myotis myotis*, *Myotis nattereri*, *Nyctalus leisleri*, *Nyctalus noctula*, *Pipistrellus nathusii*, *Pipistrellus pipistrellus*, *Pipistrellus pygmaeus*, *Plecotus auritus*, *Plecotus austriacus*, *Barbastella barbastellus* | *Vespertilionidae*: *Nyctalus leisleri* (Lesser noctule), *Plecotus auritus*, *Plecotus austriacus* (Grey long-eared bat), |
| Ireland (12 sampling sites) | 366 | *Vesperitlionidae*: *Nyctalus leisleri*, *Pipistrellus* spp., *Plecotus auritus*<br>*Rhinolophidae*: *Rhinolophus hipposideros* | *Vesperitlionidae*: *Pipistrellus pygmaeus* - Soprano pipistrelle |
| **Total** | 2667 |  |  |

Expected species: Bats species known to circulate/inhabit the area were based on prior local monitoring studies. Identified species: Bat species barcode using Cox1, cytochrome b or 16S rRNA sequencing upon coronavirus sequence detection in sample.

**Figure 1.**
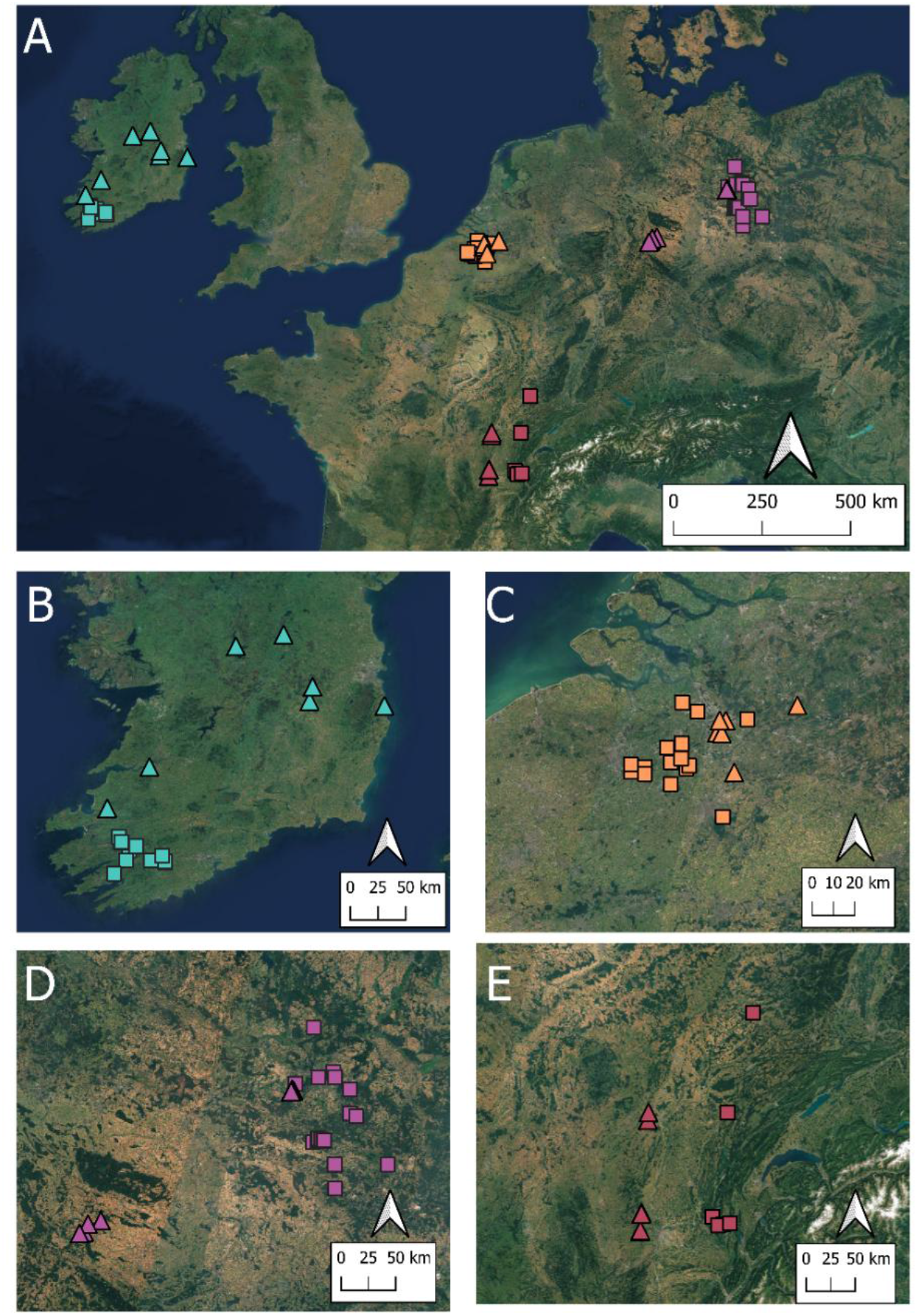
Geographical distribution of the samples analysed for coronaviruses. Terrestrial small mammals (pentagons) and bat guano (squares) samples recovered from sampling sites across four countries were analysed in this study. The distribution of samples is shown (A) across the continent and within (B) Ireland (cyan), (C) Belgium (orange) and (D) Germany (magenta), and (E) France (red). The base map is available in EU’s Copernicus Sentinel Hub, Sentinel 2 satellite images - European Space Agency (ESA). 2023. Sentinel–2 Global Mosaic (S2GM) – Basemap. Copernicus Land Monitoring Service. https://s2gm.land.copernicus.eu

### Detection of CoVs in rodent and bat samples

Based on PCRs targeting the RNA-dependent RNA polymerase (RdRp) gene, we detected coronaviruses in 59 samples in this study, which constituted 23 rodent and 36 bat samples (S1 Table). Several rodent species and shrews tested in this study were negative for coronaviruses, likely due to their low sample size, limiting our ability to detect positive individuals (S1 Table).

#### Prevalence and distribution of coronaviruses in rodents

Across the four countries, rodent coronavirus (CoV) prevalence was highest in Belgium was (4.17%; 95% CI 2.24-7.02), followed by France (1.95%; 95% CI 0.90-3.67), and Ireland (0.45%; 95% CI 0.01-2.50), whereas no positive samples were detected in Germany. Half of the sampling sites (3 out of 6) in Belgium and France had positive samples, while only one out of six sites studied in Ireland was positive (S2 Table).

We found CoVs in *A. flavicollis* (3.43%, 95% CI 1.39-6.94), *R. norvegicus* (2.94%, 95% CI 0.07-15.33), *C. glareolus* (2.11%, 95% CI 1.01-3.84) and *A. sylvaticus* (1.34%, 95% CI 0.44-3.11) (S3 Table). Additionally, *C. glareolus* tested positive for coronaviruses across multiple sampling sites, suggesting its potential importance in the circulation of these viruses (S2 Table). However, an assessment using Generalised linear models (GLMs) indicated no significant association between host species and the presence of coronaviruses in general (S4 Table, S5 Table).

The global site-specific variation was mostly small (S4 Table), even in the GLMs fitting Belgium and France separately (S5 Table). However, we found a marginally significant effect of site within countries for Belgium, particularly at the Planckendael Zoo. Estimated marginal means showed that predicted prevalence was close to zero in most sites, with higher values observed at the Planckendael Zoo (OR= 20, 95% CI = [4.33–140], *p* = <0.001).

#### Distribution of coronaviruses in bats

CoV sequences were detected in bats from all four countries, spanning three bat families: 18 from *Vespertilionidae*, 15 from *Rhinolophidae* and one from *Miniopteridae*. Previous studies have reported CoVs from *P. pygmaeus*, *P. auritus*, *N. leisleri, R. ferrumequinum*, *R. hipposideros*, and *M. schreibersii* ^5^. We confirm detections in these species and additionally report coronaviruses in two vesper bats, *P. austriacus* and *M. mystacinus*.

Coronavirus positive samples were found at three out of 16 sites in Belgium, (Halle-Zoersel, Rood Klooster and Vrieselhof), three out of 21 sites in Germany (Rochauer Heide, Doberlug-Kirchhain and Thüren), four out of the five sites in France (Poligny, Sabla, Tenay and Carroussel cave) and one out of 21 sites in Ireland (Killarney National Park). As guano was collected from the environment and could not be confidently assigned to individual bats, species identification was confirmed only for positive samples, and no statistical analyses was performed. Bat species could not be determined for three positive bat samples.

### Characterisation of CoVs detected in rodents and bats

To assess coronavirus diversity, we conducted phylogenetic analysis based on 60 RdRp sequences obtained from 59 samples through sequencing from 23 rodent and 36 bat samples. Most sequences were generated by Sanger sequencing, with an average length of 400 nt. Four samples yielded complete RdRp sequences obtained from next generation sequencing. One of these samples (PA20) contained two distinct coronavirus sequences, both of which were included in the analysis. Two sequences were obtained from bat guano pools. The analysis showed that 19 sequences clustered within the *Alphacoronavirus* genus and 41 within the *Betacoronavirus* genus (Figures 2 and 3), with representative sequences from both bats and rodents of all countries included in this study.

**Figure 2.**
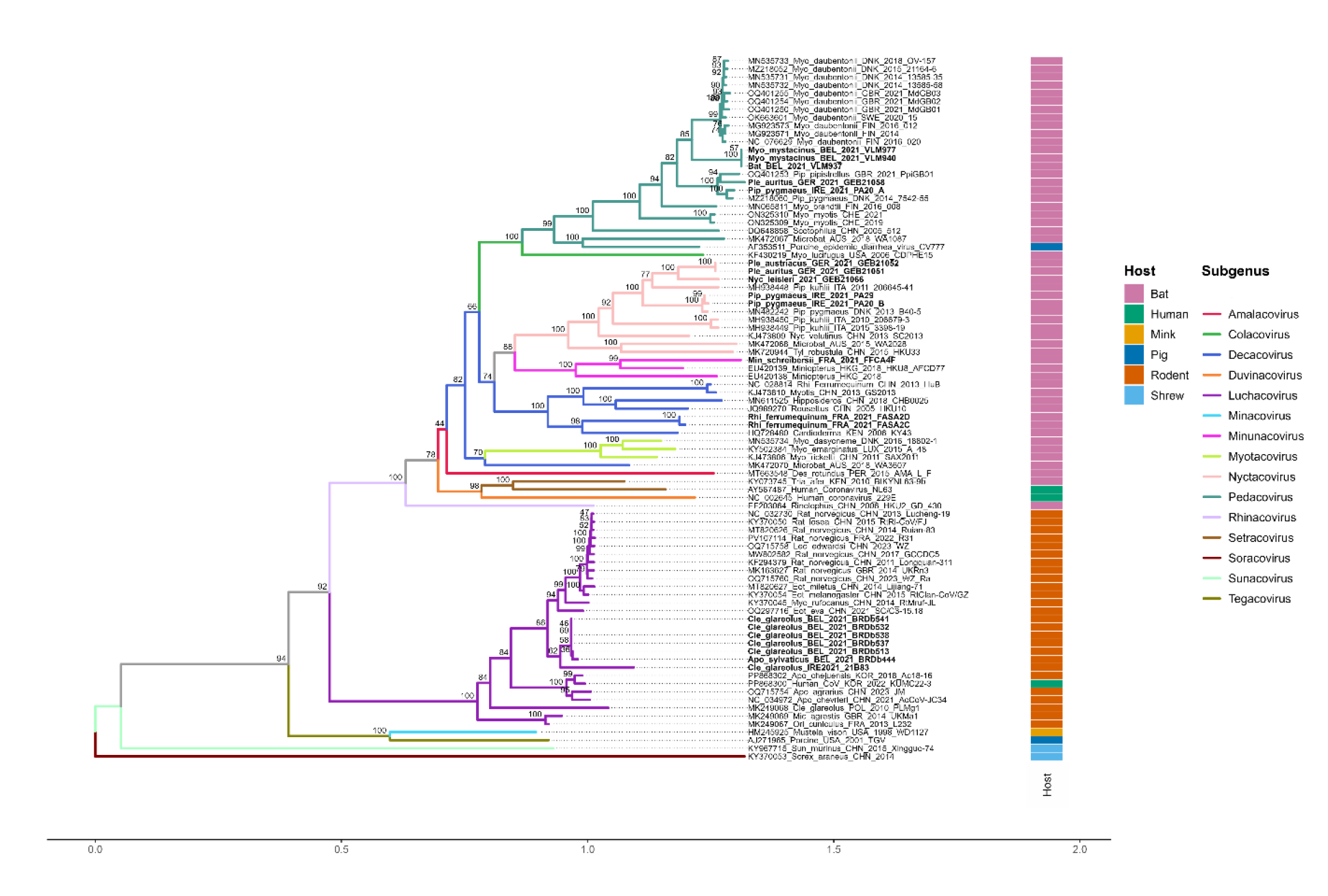
Alphacoronavirus phylogenetic tree constructed based on RdRp gene using IQ-TREE 3, with model GTR+F+I+G4), branch support assessed using 1,000 ultrafast bootstrap replicates. The final tree topology was visualised and annotated using the R studio package ggtree^33–37^. Refer to the methods section of phylogenetic analysis for details of the length of sequences analysed.

**Figure 3.**
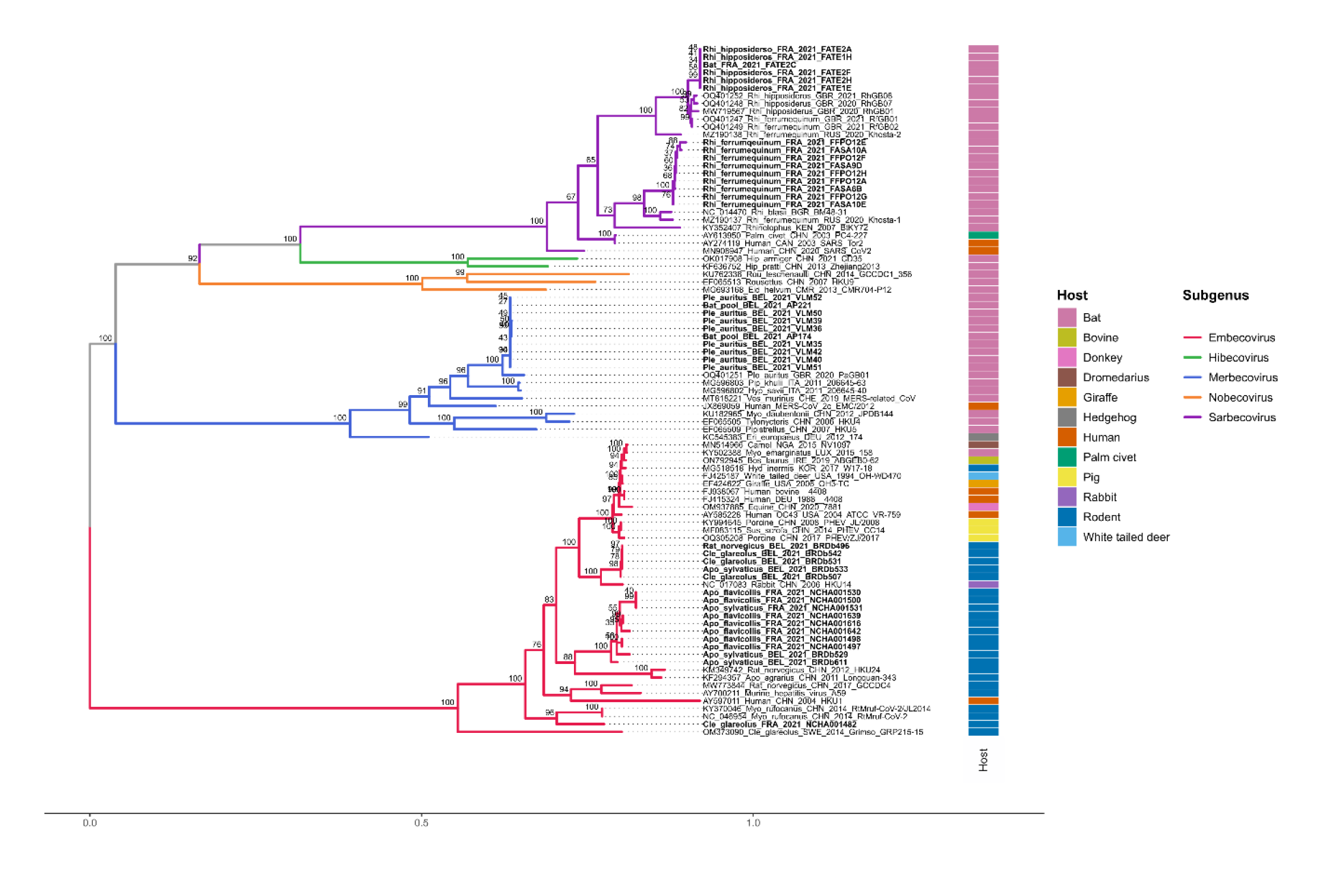
Betacoronavirus phylogenetic tree constructed based on RdRp gene using IQ-TREE 3, with model GTR+F+I+G4), branch support assessed using 1,000 ultrafast bootstrap replicates. The final tree topology was visualised and annotated using the R studio package ggtree^33–37^. Refer to the methods section of phylogenetic analysis for details of the length of sequences analysed.

#### Alpha- and Betacoronaviruses in European rodents

The phylogenetic analysis revealed that the 23 rodent-CoVs detected in this study distributed across both the *Alphacoronavirus* and *Betacoronavirus* genera, with seven clustering within genus *Alphacoronavirus* (Fig 2) and 16 within genus *Betacoronavirus* (Fig 3) (GenBank PX227390 -PX227409, PZ322830-PZ322832).

The rodent alpha-CoVs and beta-CoVs identified in this study clustered with previously characterised rodent coronaviruses of the respective genera and subgenera. Five *C. glareolus* CoV sequences from Planckendael Zoo together with one CoV sequence from *A. sylvaticus* from the same site (BRDb444) and one more sequence from *C. glareolus* sampled in Ireland, (21B83) formed a monophyletic cluster within the subgenus *Luchacovirus*. The five *C. glareolus* sequences from Belgium shared >99% nucleotide identity with each other and 96% identity with the *A. sylvaticus* sequence from Belgium. Sequence identity between the *C. glareolus* viruses from Belgium and Ireland exceeded 87%. Further analysis showed that this rodent-associated clade shared a recent common ancestor with a human coronavirus identified in Korea (PP868300) (Figure 2, S1 Figure, S9 Table)^38^.

The identified rodent betacoronaviruses formed three distinct clades within the subgenus *Embecovirus* (Figure 3, S2 Figure, S10 Table). One clade comprised of viruses from five rodents captured at Planckendael Zoo, Belgium, including three *C. glareolus*, one *R. norvegicus*,and one *A. sylvaticus*. The *C. glareolus* and *R. norvegicus* sequences shared >99.7% nucleotide identity, whereas the *A. sylvaticus* sequence shared <90% identity with the other viruses of the clade.

The second clade consisted of 10 beta-CoV sequences detected in seven *A. flavicollis* and three *A. sylvaticus* from Belgium and France, including samples from Planckendael Zoo and Zevendonk in Belgium, and Cormaranche, Griffe au Diable, and Mignovillard in France. Despite the large geographic distances (about 94 kms apart) separating these sampling locations, the sequences shared 95–98% nucleotide identity.

Finally, a single beta-CoV sequence detected in *C. glareolus* from Griffe au Diable, France clustered separately from the previously described clades. This virus clustered with rodent coronavirus RtMruf-2 (NC_046954), originally identified in *M. rufocanus* in China, sharing 88.2% nucleotide identity with it. Although distinct from the other *Embecovirus* clades, this virus shared a common ancestor with them and exhibited 75–83% nucleotide identity with the other rodent *Embecovirus* sequences identified in this study (Figure 3, S2 Figure, S10 Table).

Post virus characterisation, statistical analyses using Fischer’s Exact and Pearson’s Chi-sqaure tests were performed to understand the association between host species and the genera of coronaviruses they carried. We found a positive association of Alphacoronaviruses to *C. glareolus* and Betacoronaviruses to *Apodemus species* (S11 Table). Also, the same tests were performed to check association of coronavirus genera detected by country, and we found an association of Alphacoronaviruses to Belgium and Betacoronaviruses to both Belgium and France (S11 Table). Both these results corroborate our findings from the prevalence studies and phylogenetic analyses.

#### Alpha- and Betacoronaviruses in European bats

Phylogenetic analyses showed that the identified bat CoVs were also distributed across the *Alpha*- and *Betacoronavirus* genera (GenBank PX2273953 - PX227389, PZ322830 - PZ322833, PZ723670). The bat alpha-CoVs clustered within subgenera *Nyctacovirus*, *Decacovirus*, *Minunacovirus* and *Pedacovirus*, whereas the bat beta-CoVs clustered within subgenera *Merbecovirus* and *Sarbecovirus*.

Two sequences from *R. ferrumequinum* collected in Sabla, France, grouped within *Alphacoronavirus* subgenus *Decacovirus*, sharing 98.87% nucleotide similarity with one another.

Within the *Alphacoronavirus* subgenus *Nyctacovirus*, sequences from *P. austriacus* and *P. auritus* were identical to one another. Together these formed a sister clade to *Nyctalus leisleri* sequence from Germany, with 88.45% nucleotide level identity. Two sequences from *P. pygmaeus* from Ireland, sharing 98.3% identity within themselves, clustered in a separate clade. All these sequences clustered in sister lineages, indicating a shared evolutionary origin (Figure 2, S1 Figure, S9 Table). A single sequence from an *M. schreibersii* of France clustered within *Alphacoronavirus* subgenus *Minunacovirus*, together with other coronaviruses isolated from *Miniopterus* bats.

Within the subgenus *Pedacovirus*, two *M. mystacinus* sequences and one sequence from an unidentified bat from Belgium (VLM937) formed a distinct cluster and shared 100% nucleotide identity. Additionally, two sequences - one from *P. auritus* in Germany and the other from a *P. pygmaeus* in Ireland - shared 93.4% identity and were placed in sister clades within the subgenus *Pedacovirus*. These pedacoviruses though share a second-generation ancestor.

Two different alphacoronaviruses were identified in one Irish *P. pygmaeus* sample (PA20), where one of the CoV sequences (PA20_A) belongs to the *Pedacovirus* subgenus, while the other (PA20_B) belonged to the *Nyctacovirus* subgenus.

Bat beta-CoV sequences were distributed in three clusters, within the subgenera *Sarbecovirus* and *Merbecovirus*. Eight sequences detected from bats captured in Belgium, *P. auritus* and two from *P. auritus* pools (AP174, AP221), were nearly identical, and formed one distinct lineage within the *Merbecovirus* subgenus (Figure 3, S2 Figure, S10 Table). These viruses share a distant ancestor with MERS-CoV-2.

The sequences obtained from *Rhinolophus* bats in France formed two distinct clusters under the *Sarbecovirus* subgenus. The first cluster contained five sequences from *R. hipposideros* and one sequence from an unidentified bat (FATE2C), were almost identical to one another. The second cluster of nine sequences from *R. ferrumequinum* shared 97% to 100% sequence identity with one another. These sequences share a distantly related to SARS-CoV and SARS-CoV-2.

It was noted that the merbecoviruses from this study were found only in *Plecotus* bats and the sarbecoviruses in *Rhinolophus* bats, specifically *R. ferrumequinum and R. hipposideros*.

## Discussion

In this study, we demonstrate the diversity of coronaviruses circulating in rodents and bats across four European countries during Spring 2021. Among the countries surveyed, Belgium exhibited the highest coronavirus prevalence, exceeding prevalence reported in previous studies of European rodents and bats^13–15,39–42^. Together, our findings suggest substantial regional heterogeneity in coronavirus circulation.

At the local scale, most rodent coronaviruses identified in this study were in Belgium and France. In Belgium, nearly all positive samples were collected from Planckendael Zoo, with only two additional positives detected at Fort 5 and Zevendonk. Rodents sampled from Belgium carried both alphacoronaviruses and betacoronaviruses, indicating the co-circulation of multiple coronavirus genera within the same ecological setting. Positive animals belonged to several species, including *A. sylvaticus*, *C. glareolus*, and *R. norvegicus*, with *C. glareolus* accounting for most of the detections. The simultaneous circulation of alpha- and betacoronaviruses in sympatric rodent populations may increase opportunities for cross-species transmission and recombination, both of which are recognised as important drivers of coronavirus evolution^43–46^.

Phylogenetically, the alphacoronaviruses detected in Belgium clustered within the subgenus *Luchacovirus*. Notably, the viruses identified in *C. glareolus* from Planckendael Zoo formed a distinct cluster separate from previously described European luchacoviruses. Earlier reports identified related viruses primarily in rats and voles from the United Kingdom, France and China^41,47,48^. Our findings therefore expand the known host range of this lineage and its geographical location within Europe and suggest additional genetic diversity within the subgenus. Interestingly, these viruses were also distantly related to a recently described rodent alphacoronavirus associated with human infections in South Korea^38^, underscoring the link between rodent and human coronaviruses and patterns of possible zoonotic events.

The rodent betacoronaviruses detected in Planckendael Zoo belonged to the subgenus *Embecovirus*. Phylogenetic analysis revealed three distinct lineages: two major clusters and a single divergent sequence forming an independent branch. One cluster consisted exclusively of sequences from Belgium, whereas the second contained mainly French sequences together with some sequences from Planckendael Zoo and Zevendonk. *Embecoviruses* are notable for their broad host ranges, a characteristic that was also evident in our dataset where the Belgian lineage was identified in *C. glareolus*, *R. norvegicus*, and *A. sylvaticus*, and the French lineage was detected in both *A. sylvaticus* and *A. flavicollis*. Beyond rodents, embecoviruses have been reported in a wide range of mammalian hosts, including captive zoo animals^49,50^. These observations highlight the ecological flexibility of embecoviruses and support concerns that free-ranging rodents may facilitate viral exchange between wildlife and captive animal populations. As rodents frequently move between natural habitats and zoological enclosures, they may contribute to the introduction, maintenance, or spread of coronaviruses within zoo environments, emphasising the importance of targeted surveillance at this interface^51^.

Taken together, the detection of both alpha- and betacoronaviruses in rodents from Planckendael Zoo highlights zoological parks as environments where diverse coronavirus lineages may co-circulate within a relatively confined area. Previous studies have documented pathogen movement between free-ranging rodents, birds and zoo animals^46,52,53^. These interactions can occur directly or indirectly through shared food resources, enclosure surroundings or transport-associated activities^13,19^. Because captive animals may be particularly susceptible to pathogen introduction from surrounding wildlife, surveillance of wildlife inhabiting zoos, alongside effective pest management strategies, represents an important component of zoo biosecurity. Beyond the implications for animal health, zoos are also notable as interfaces where wildlife, captive animals and humans come into close contact, creating conditions that may facilitate pathogen exchange across taxonomic boundaries^53,54^.

In contrast to Belgium, all rodent coronaviruses detected in France belonged to the subgenus *Embecovirus*. Positive animals were restricted to *A. sylvaticus* and *A. flavicollis*, and the sequences identified in France formed a distinct phylogenetic cluster separate from previously described rodent coronaviruses in France^14,41^. A similar observation was made for the Belgian luchacoviruses and one of the embecovirus clusters identified in this study, both of which occupied phylogenetically distinct positions relative to previously reported viruses. Collectively, these findings suggest the presence of previously uncharacterised coronavirus diversity circulating within rodent populations in Belgium and France. Although most detections were based on partial RdRp fragments, two complete RdRp sequences generated through next-generation sequencing (BRDb538 within luchacoviruses and NCHA001531 within embecoviruses) also formed divergent lineages relative to currently available reference sequences, providing additional support for this interpretation.

Evidence from our study also suggest host-associated circulation patterns among the rodent coronaviruses. Embecoviruses in France were identified exclusively in *Apodemus* species, whereas the luchacoviruses from Belgium were associated with *C. glareolus*, which was also the dominant coronavirus-positive species within the Planckendael Zoo. Statistical analyses further showed associations between *C. glareolus* and alphacoronavirus detection and between *Apodemus* and *Clethrionomys* species and betacoronavirus detection. These observations are consistent with the possibility of host-associated circulation patterns, particularly for the luchacoviruses. However, given the limited number of positive samples and the known occurrence of rodent alphacoronaviruses in multiple host species across Europe, these association patterns should be interpreted cautiously. Larger studies incorporating genomic and co-evolutionary analyses will be required to determine whether these patterns reflect true host specificity or ecological sampling effects^13,14,55^.

Similarly, the single positive rodent detected in Ireland was identified as *C. glareolus*, and its coronavirus clustered with the other luchacoviruses identified in Plackendael Zoo, Belgium. Given that the Irish bank vole population is believed to have originated from a small founder population following introduction to Ireland in the 1920’s^56^, the occurrence of a closely related virus lineage raises interesting questions regarding host-associated maintenance and historical virus dispersal across Europe^57^. Although the small sample size limits our ability to draw definitive conclusions, this observation is consistent with the hypothesis of potential association between *C. glareolus* and this alphacoronavirus lineage discussed above. Ireland’s geographic isolation as an island nation has resulted in lower rodent species diversity than mainland Europe, which may contribute to reduced pathogen diversity and fewer opportunities for viral diversification through host switching or recombination^58,59^.

Contrary to our expectation, no coronavirus-positive rodents were detected in Germany despite previous reports of rodent coronaviruses from the country^13^. However, the lack of positive samples does not rule out the circulation of viruses in the surveyed areas. Methodological differences may partly explain this discrepancy. While samples from the other study countries were collected from live-captured animals, German samples were obtained from animals recovered dead from modified traps, with tissues collected up to 24 hours after capture. Such delays might have compromised RNA quality and reduced detection sensitivity^60^. Moreover, ecological factors such as seasonality, population density, population behaviour, sympatric populations and local host movement patterns may also have influenced detection probability^61^.

Our study primarily sampled voles, mice and rats, a bias based on traps and bait used for this study. Although a small number of shrews and squirrels were also captured, no coronaviruses were detected in these species. However, coronaviruses have previously been reported from a range of European small mammals, including hedgehogs, rabbits, squirrels and shrews^13,14,25,26,62^. Given the limited number of non-rodent small mammals sampled, our study could not assess coronavirus diversity across all potential hosts. Furthermore, geographical coverage of this study was limited to the patchy sample sites located within four European countries only. Future studies incorporating a broader range of host species, sampling locations and countries, would provide a more complete understanding of coronavirus diversity and host associations across Europe. Such efforts may be particularly valuable for examining differences in coronavirus diversity between island populations and continental wildlife communities.

In contrast to the rodent findings, the bat coronaviruses detected in this study largely reflected previously recognised host-virus associations. An alphacoronavirus belonging to the subgenus *Minunacovirus* was identified in a French *Miniopterus schreibersii* bat (FFCA4F) sampled from a forest in Mignovillard and clustered with previously described minunacoviruses. This observation is consistent with previous studies demonstrating a strong association between minunacoviruses and *Miniopterus* bats^63^. Similarly, coronaviruses detected in *R. hipposideros* and *R. ferrumequinum* from France clustered within the subgenus *Sarbecovirus* alongside previously reported Rhinolophus-associated sarbecoviruses. Together, these findings reinforce the growing body of evidence indicating that several bat coronavirus lineages exhibit long-term evolutionary associations with specific bat taxa^1,8,64^. Consistent with previous reports, the sarbecoviruses identified in *Rhinolophus* bats from Europe were phylogenetically distant from both SARS-CoV and SARS-CoV-2^7,8,22,23,64–66^. Likewise, the merbecoviruses detected in *Plecotus auritus* remained distantly related to MERS-CoV.

Beyond host associations, we identified evidence of coronavirus co-circulation in bats at an individual level. Two alphacoronaviruses belonging to different subgenera were detected in a *P. pygmaues* from Ireland, including PA20_A from the subgenus *Pedacovirus* and PA20_B from the subgenus *Nyctacovirus*. This finding demonstrates that individual bats can harbour multiple coronavirus lineages simultaneously and is consistent with previous reports of coronavirus co-infections in bat populations^64,65^. Co-circulation, particularly of alpha- and betacoronaviruses, is evolutionarily significant because it can promote viral recombination, a key mechanism underlying coronavirus diversification and host adaptation^7,8^.

Interestingly, the merbecoviruses identified in bats from Belgium formed a monophyletic cluster and were most closely related to previously detected merbecoviruses from the United Kingdom. This pattern may reflect localised evolution of merbecoviruses within *Plecotus* bat populations. Such a scenario is biologically plausible given the ecology of *Plecotus* bats, which generally exhibit limited dispersal distances and do not frequently co-roost with other bat species^7,64,67^. Restricted movement and relatively closed host populations may promote long-term virus-host co-evolution, resulting in geographically structured viral lineages as observed here.

Despite the limitations associated with sampling coverage, host representation and variation in sampling protocols among countries, this study expands our understanding of coronavirus diversity in European terrestrial small mammals and bats. While the bat coronaviruses largely conformed to previously recognised host-associated lineages, rodent sampling revealed substantial diversity. In particular, the detection of both alpha- and betacoronaviruses among rodents inhabiting Planckendael Zoo highlights the value of zoological collections as One Health surveillance sites where pathogen transmission dynamics can be investigated across wildlife, animal and human interfaces. Although the coronaviruses detected in this study do not indicate an immediate zoonotic threat, their circulation within such interfaces underscores the importance of continued surveillance to better understand viral diversity, transmission dynamics and potential future host-switching events.

Future studies incorporating complete genome sequencing, particularly of the spike protein, would enable more detailed investigations of receptor usage, host adaptation and recombination patterns. Such analyses are especially important because the emergence of human coronaviruses typically reflects prolonged evolutionary processes involving adaptation in animal hosts and, frequently, transmission through intermediate hosts rather than direct spillover events^68^. Understanding the genomic features that facilitate host switching will therefore be critical for assessing the emergence potential of newly identified wildlife coronaviruses.

## Materials and Methods

### Ethics statement

All procedures complied with relevant national, European, and institutional guidelines for the use of animals in research, including Directive 2010/63/EC revising Directive 86/609/EEC. In Germany, rodent trapping, handling, and sampling were conducted under the permits of Landesamt für Arbeitsschutz, Verbraucherschutz und Gesundheit Brandenburg (LAVG no. 2347-A-16-1-2020), Landesamt für Umwelt Brandenburg (LfU no. LFU-N1-4744/97+17#194297/2020), and by permission of the Thuringian State Office of Consumer Protection (TLV; permit no. 22-2684-04-15-105/16). In France, the CBGP laboratory is authorised by the Departmental Direction of Population Protection (DDPP, Hérault; approval F-34-169-001) for the sampling of small mammals and the storage and use of their tissues. Procedures were validated by the regional ethics committee “Comité d’Éthique pour l’Expérimentation Animale Languedoc Roussillon n°36” in 2020 (ref: 2020-02-v2). In Ireland, rodent sampling was approved by the Munster Technological University ethics board and was conducted in compliance with the Health Products Regulatory Authority (HPRA) for euthanasia of sampled rodents (Authorisation AE22171/I004).

Bat faecal samples in Germany and France were collected from artificial roosts accessed during routine maintenance activities that did not require permits. In Belgium, all bat sampling was approved by the ethical commission of the University of Antwerp and adhered to regulations of the Belgian Agency for Nature and Forest (ANB), including applicable COVID-19 measures at the time of sampling. In Ireland, bat surveys were conducted in compliance with the Irish National Parks and Wildlife Service under survey licence DER/BAT 2021-87.

### Sample collection and storage

The terrestrial small mammal samples were collected as part of the Biodiversa/BioRodDis project (https://www.biodiversa.eu/2022/10/31/bioroddis/)^69,70^. The focus of this study was on the subset of terrestrial small mammal samples collected from March to June 2021 in France, Belgium, Germany and Ireland (Figure 1, Table 1, S7 Table). The shrews sampled were found dead in the traps in France and Belgium. In Ireland and Germany, modified traps^71^ were used to avoid shrew trapping. The animals were morphologically identified in the field, and whenever necessary by molecular analysis. Sections of colon were collected in RNAlater (Invitrogen MA, USA) and stored at -20 °C for coronavirus RNA preservation and detection.

Bat guanos were collected in France, Belgium, Ireland and Germany from March to September 2021 as detailed S7 Table (Figure 1, Table 2; and details in S8 Table)^72^. Sampling sites included a natural cave in Ireland; forested areas in France; and buildings such as churches and historical landmarks in Belgium and Ireland. While the samples from France, Belgium and Ireland were collected from under roosts, German samples were collected from the floor underneath colonies roosting in private buildings and from inside bat boxes.

A clean, fresh, white paper sheet was put on the floor under a bat colony or where a bat colony was expected based on the fresh dropping found on the floor. The freshest faecal pellets were collected the following day and put separately in Eppendorf tubes filled with RNAlater. After 24 hours in a fridge (4 °C), the guano pellets were stored in RNAlater and -20 °C until further use. The expected bat species at the sampling sites were identified via acoustics monitoring in France and based on prior knowledge in Belgium, Germany and Ireland^27–32^.

### Bat identification

Only CoV positive samples were barcoded to identify bat species. Bats from Belgium were identified using cytochrome b (cyt b) and 16S rRNA sequencing at the Neuromics Support Facility of the Vlaams Instituut voor Biotechnologie (VIB) in Antwerp, Belgium. The bats from France and Ireland were identified through cytochrome c oxidase I (COI) gene sequencing.

A short region of mitochondrial cytochrome b covering 476bp was amplified using the protocol in S5 Table, Section E and Supplementary Material 1, and sequenced using the L14723_Cytb forward and H15149_short reverse primers as per previous publications^73–75^. For the samples which failed Cytochrome b sequencing, 16S rRNA gene was sequenced instead using the 16Smam1 and 16Smam2 primers targeting a 130bp fragment as per prior studies^76^. For the bats from France and Ireland, the mitochondrial cytochrome C oxidase subunit I (COI) was sequenced for regions spanning 133 bp and 185 bp respectively using MG2-LCO1490 forward and MG2-univ-R primers as in previous publications^77^.

### Samples preparation for RNA extraction

For RNA extraction we used the QIAamp 96 Virus QIAcube HT kit and QIACube HT extraction robot giving a final RNA elution volume of 120µl (QIAgen, Germany). Total RNA extracted was stored at -80 °C for downstream processing, and the homogenates were stored at -20°C.

Colon samples (5 mg) were processed by enzymatic and mechanical lysis to produce a digested homogenate, which was then clarified by centrifugation (12000 G for 1 minute) and used for RNA extraction using the off-board lysis protocol according to manufacturer’s instruction. Detailed lysis protocol can be found in Supplementary Material 2.

Bat guano samples were washed from RNAlater prior homogenization. The guano lysate was produced by mechanical homogenization using QIAGEN TissueLyser II (QIAgen, Germany) and clarified by centrifugation (12000 G for 1 minute). A total of 200 ul of clarified lysate was used for RNA extraction using onboard lysis protocol following the manufacturer’s instructions. Lysate samples from Belgium were pooled in groups of 7-9 individuals before RNA extraction. The other samples were processed individually (S3 Figure). Detailed information on how the bat guano were processed for RNA extraction can be found in Supplementary Material 3.

### Coronavirus screening and confirmatory PCRs

The screening PCR was done using methods adapted from and primers described in Escutenaire et. al. (2007)^78^. This PCR was performed using the AriaMx Real-time PCR System (Agilent Technologies, Santa Clara, CA, USA) and the data generated was analysed using the AriaMx Software (Agilent Technologies, Santa Clara, CA, USA). Detailed protocol can be found in S5 Table, Section A.

Maxima Reverse Transcriptase (Thermo Scientific™) was used in combination with RiboLock RNase Inhibitor (Thermo Scientific™) and random primers in reactions containing 5 ul of total RNA to produce cDNA. Detailed methodology on the cDNA synthesis can be found in S5 Table, Section D. Prepared cDNA was stored at -20 °C until further use for PCR and sequencing. Confirmatory semi-nested PCR A was performed by modifying the protocol of and utilising primers from Hoolbrook et. al. (2021)^79^. The confirmatory nested PCR B was performed by modifying the protocol and utilising the using the primers from de Souza Luna et al. (2007)^80^. More details on the PCRs can be found in S6 Table Sections B and C.

Positive amplicons from the conventional PCRs were sent for final confirmation via amplicon sequencing. Amplicons of expected length (420-450 bp) were purified using GeneJet PCR purification kit (ThermoScientific, MA, USA) or GeneJet Gel extraction kit (ThermoScientific, MA, USA) as per manufacturer’s instructions. Amplicon sequencing was performed at the FIMM sequencing facility (University of Helsinki) using Applied Biosystems BigDye Terminator v3.1 Cycle Sequencing Kit (Part No. 4336921) and analysed with Applied Biosystems ABI3130XL Genetic Analyzer (16-capillaries), following the recommended protocols.

### Next generation sequencing and analysis

Two coronavirus-positive samples each from rodents and bats were selected for next generation sequencing (NGS), to derive complete or near complete CoV RdRp sequences of representative Alpha-CoVs and Beta-CoVs. Samples from RNA extracted from rodent colon (NCHA001531 and BRDb538) for the PCRs were used for sequencing. Bat faeces (PA20 and FASA10E) were processed following the NetoVIR protocol^81^ and total nucleic acids extracted manually using the QIAamp Viral RNA Mini Kit (Qiagen, MA, USA). cDNA synthesis and amplification were carried out using the Complete Whole Transcriptome Amplification II Kit (Sigma-Aldrich, MI, USA). Sequencing libraries were prepared using the NEBNext® Ultra™ II RNA Library Prep Kit for Illumina® (New England Biolabs, MA, USA). dsDNA fragmentation was done using NEBNext® Ultra™ II FS DNA Library Prep Kit for Illumina (New England Biolabs, MA, USA) to obtain fragments of 100 – 250 bp. All libraries were sequenced with the Illumina NovaSeq 6000 system at the FIMM Sequencing Facility, FIMM Genomics Services, University of Helsinki, Finland. The raw sequencing reads were processed using LazyPipe v3.1^82,83^, a de novo assembly bioinformatic pipeline for virus discovery. The annotation strategy was recommended for Illumina libraries from environmental and faecal samples with complex eukaryotic backgrounds^84^ (S 12 Table). Briefly, Minimap2^85^ was used to query the de novo assembled contigs against the minimap.nt database, followed by a DIAMOND^86^ search against the uniref100.abv database for all contigs that remained unassigned.

### Phylogenetic analysis and similarity matrix

A total of 166 RdRp gene sequences, including 110 CoV reference gene sequences retrieved from GenBank, both complete and partial, were used for phylogenetic analysis. The dataset comprised predominantly partial RdRp sequences amplified by PCR and Sanger sequencing, together with four complete RdRp coding sequences (CDS) obtained by next-generation sequencing (NGS) from samples NCHA001531, BRDb538, PA20 and FASA10E.

Sequence alignments were generated using MAFFT version 7, with the E-INS-I refinement method^87^. Maximum-likelihood phylogenies were inferred using IQ-TREE3, incorporating ModelFinder to determine the best-fitting nucleotide substitution model^88,89^. Branch support was assessed using 1,000 ultrafast bootstrap replicates following previously described methods^89^. Based on the Bayesian Information Criterion (BIC), ModelFinder selected GTR+F+I+G4 as the best fitting substitution model. The final tree topology was visualised and annotated in R studio using the ggtree package^33–36^.

A pairwise sequence identity matrix was generated in RStudio using R (v4.5.2) by calculating pairwise nucleotide identities from the multiple sequence alignment while excluding gap positions. The resulting similarity matrix was visualised as a heatmap using the ggplot2 package^90^.

A tanglegram was constructed comparing the complete RdRp sequences obtained by NGS with the short RdRp sequences from Sanger sequencing. Details of methods and results can be found in Supplementary Material 4 and S4 Figure.

## Statistical analysis

All statistical tests were performed with R v4.5.2.

Positivity rate and confidence intervals for the rodent samples were calculated using base function binom.test. A generalised linear model (GLM) was used to compare species, site and country-based coronavirus detections. Due to the low prevalence of coronaviruses in our dataset, we first fitted a generalised linear mixed model (GLMM) with a binomial distribution using glmer function^91^. Species and country were included as fixed effects and site as a random effect. Model diagnostics were assessed using DHARMA package, and marginal means were estimated with the emmeans package^92,93^. This GLMM failed IN converge due to extremely low prevalence and high sparsity. We therefore performed a GLM with a nested structure (sites within countries), using a Firth logistic regression (function logistf)^94^. Consequently, we fitted a separate generalised linear model (GLM) for France and Belgium, with species and sites as fixed effects, using the glm function with a binomial distribution^95^. We investigated potential site-specific differences within each country. Fisher’s exact test with base function t.test and Pearson’s chi-squared test with base function chisq.test were performed to test for associations between coronavirus type and detection in rodent species, rodent genus, sampling site, locality and country.

## Data availability

Quality-controlled sequence data are available via the NCBI Short Read Archive (SRA), under BioProject PRJNA1456926. Sanger sequencing data of the RdRp fragments are available via NCBI GeneBank (PX227353-PX227409, PZ322830-PZ322833, PZ723670) and NGS data are available under BioProject PRJNA1456926. Refer to S13 Table for detailed information on GeneBank submissions.

## Author contributions

Author contributions are provided based on the CRediT (Contributor Roles Taxonomy) author statement. Conceptualization: ES, VV, EA, HA, VB, GC, NC, TSi, LD; Data Curation: ES, VV, VS, TSm, LD; Formal analysis: ES, VV, EA, DG, ME, MG, SGry, LD; Funding acquisition: JAE, CI, PS, LDB, GC, NC, TSi, LD; Investigation: ES, VV, EA, DG, ME, VL, TN, SGre, AV, JF, JAE, VC, AM, PS, SGry, VS, HA, VB, TSm, LD; Methodology: ES, VV, LDB, VB, GC, TSi, LD; Project administration: VV, JAE, CI, GC, NC, TSi, LD; Resources: DG, JF, JAE, JJ, CI, VS, VC, HL, AM, PS, SGry, LDB, CCV, UH, CI, TSm, GC, NC, TSi; Supervision: JAE, GC, NS, TSi, LD; Validation: ES, VV, EA, ME, TN, SGre, CI, GC, LD; Visualization: ES, VV, GC, LD; Writing – original draft: ES, VV, EA, VB, TSm, GC, NC, TSi, LD; Writing – review & editing: ES, VV, DG, ME, VL, TN, SGre, AV, MG, AM, PS, JF, JAE, JJ, CI, VS, SGry, LDB, CCV, UH, VB, TSm, GC, NC, TSi, LD. All authors read and approved the final draft, agreed to submit the manuscript and take full responsibility for its content.

## Acknowledgments

We are grateful to the people from France who collected the French Bat guano samples: Marie Parachout and Carole Simon (CPEPESC Franche-Comté for the sites Caroussel cave and Poligny), Lucie Defernez and Roxane Baudard (LPO Ain for the sites Tenay, Mussignin, Rossillon and Sabla cave). We also thank Laura McCarthy for collecting the Irish bat guano samples and to the NPWS staff and Conor Kelleher for sharing knowledge on roosts. For Flanders, we thank Robbert Schepers (Regionaal Landschap Schelde-Durme) for providing bat colony locations used in site selection and the site owners who granted access to sampling locations. The guano species identifications used in this work were partly produced at the GenSeq platform through the genotyping and sequencing facilities of ISEM (Institut des Sciences de l’Evolution-Montpellier) and analyzed thanks to the Genotoul bioinformatics platform in Toulouse Midi-Pyrénées (Bioinfo Genotoul). Lastly, we thank Viktor Olander for his technical assistance in the NGS sample preparation.

We would like to acknowledge the use of BioRender for designing the workflow illustration, the use of Microsoft Co-pilot and ChatUIT for clarity, language editing assistance and code debugging, and ResearchRabbit for reference management. LD performed bioinformatic analysis on resources provided by Sigma2 - the National Infrastructure for High-Performance Computing and Data Storage in Norway. ES and VV used CS -IT Center for Science’s Supercomputer and Data Storage in Finland for bioinformatic analysis and data storage.

## Abbreviations

PCR: Polymerase Chain Reaction
CoV/s: Coronavirus/es
hCoV: Human Coronavirus
SARS: Severe Acute Respiratory Syndrome
MERS: Middle Eastern Respiratory Syndrome
SARS-CoV-2: Severe Acute Respiratory Syndrome coronavirus 2
16s rRNA: 16s-unit ribosomal ribonucleic acid
RNA: Ribonucleic acid
RdRp: RNA-dependant RNA polymerase
GLM: Generalised linear model
OR: Odds ratio
CI: Confidence interval
p: p-value
χ²: Chi-square test coefficient
df: Degrees of freedom
W: Kendall’s W ranking
SYBR: SYBR Green I-an asymmetric cyanide dye
NGS: Next Generation Sequencing
Cyt: b Cytochrome B
COX: I Cytochrome C oxidase subunit I
bp: Base pairs
RNAlater: Ribonucleic acid later (transport media)
cDNA: Complementary deoxyribonucleic acid
TE: buffer Tris-Ethylenediaminetetraacetic acid buffer
nM: Nanomolar
mM: Millimolar
µl: Microliter
BIC: Bayesian information criterion
°C: Degree Celsius
MgCl_2_: Magnesium chloride
kHz: Kilo Hertz
CDS: Coding sequence

