## Supplementary figures for "Detection and characterisation of alpha- and betacoronaviruses in rodents and bats from Germany, France, Belgium and Ireland"

**S1 Figure: Alphacoronavirus p-distance matrix heatmap**

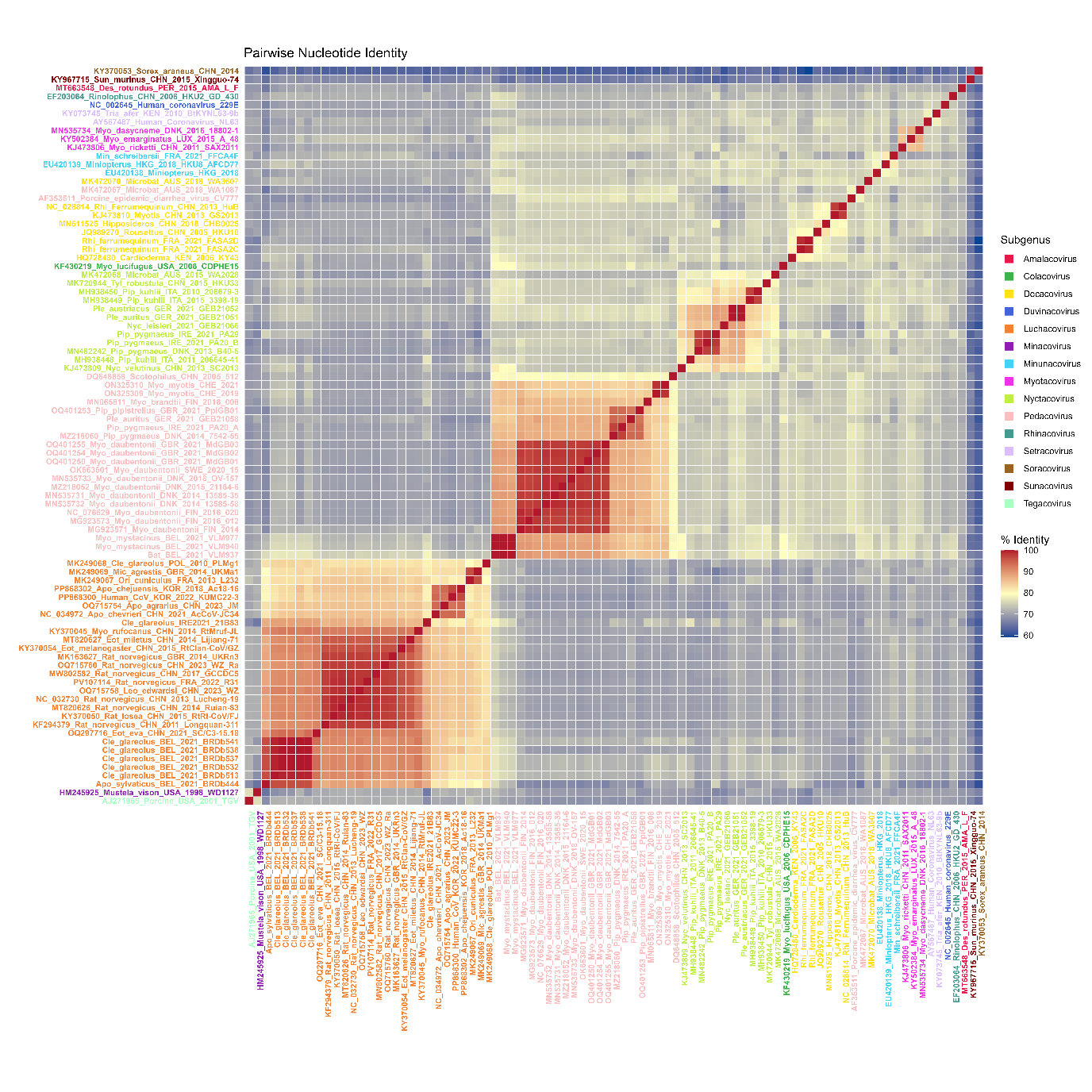

Pairwise nucleotide identity heatmap of the selected Alphacoronavirus sequences, generated using a custom R script, derivative from Table S8. Pairwise nucleotide identities were calculated from the aligned sequences after excluding gap positions, and sequence labels were color-coded according to subgenus. Colours represent nucleotide identity, ranging from lower identity (blue) to higher identity (red).

**S2 Figure: Betacoronavirus p-distance matrix heatmap**

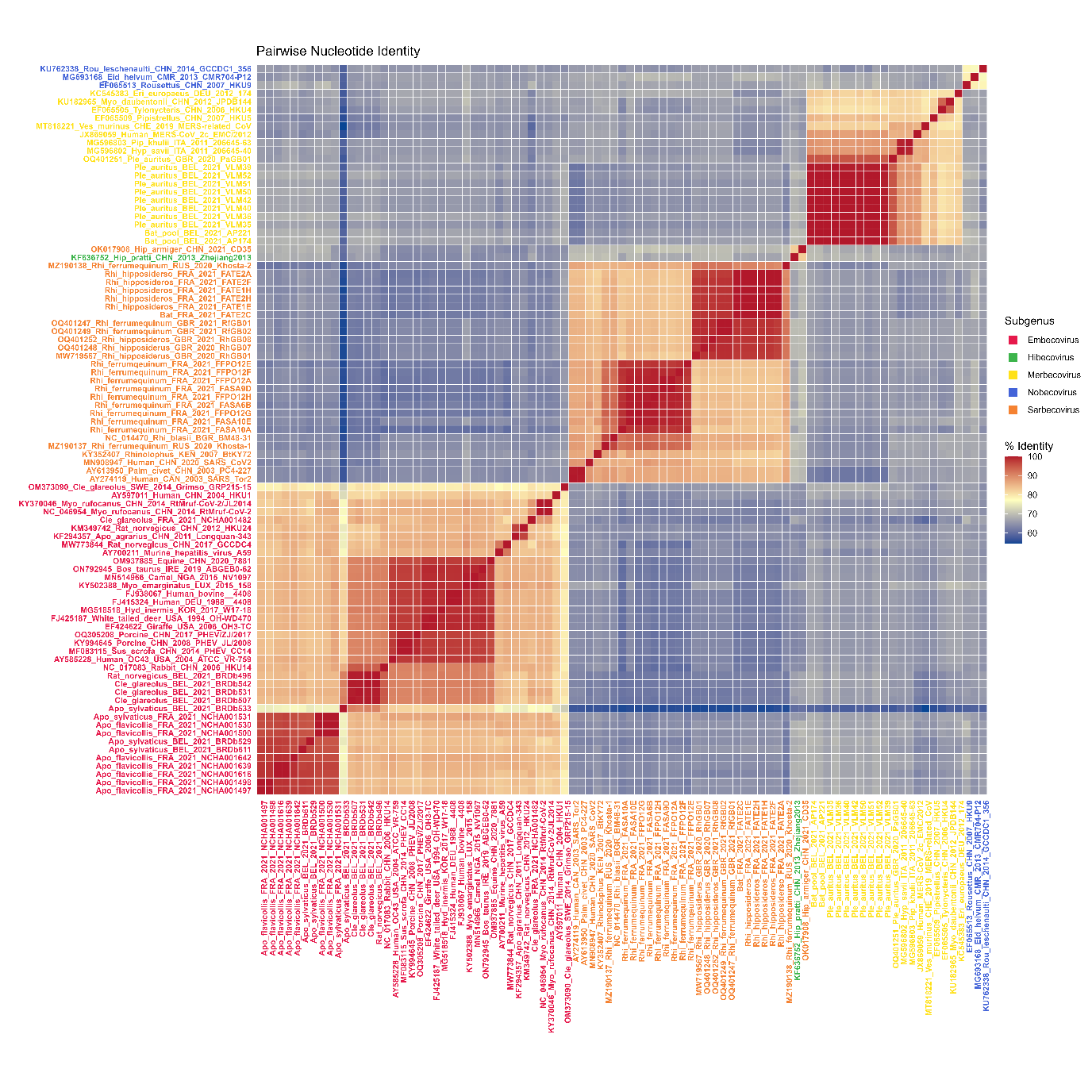

Pairwise nucleotide identity heatmap of the selected Betacoronavirus sequences, generated using a custom R script, derivative from Table S9. Pairwise nucleotide identities were calculated from the aligned sequences after excluding gap positions, and sequence labels were color-coded according to subgenus. Colours represent nucleotide identity, ranging from lower identity (blue) to higher identity (red).

**S3 Figure: Workflow for identification of coronaviruses from bat and terrestrial small mammals from all four countries.**

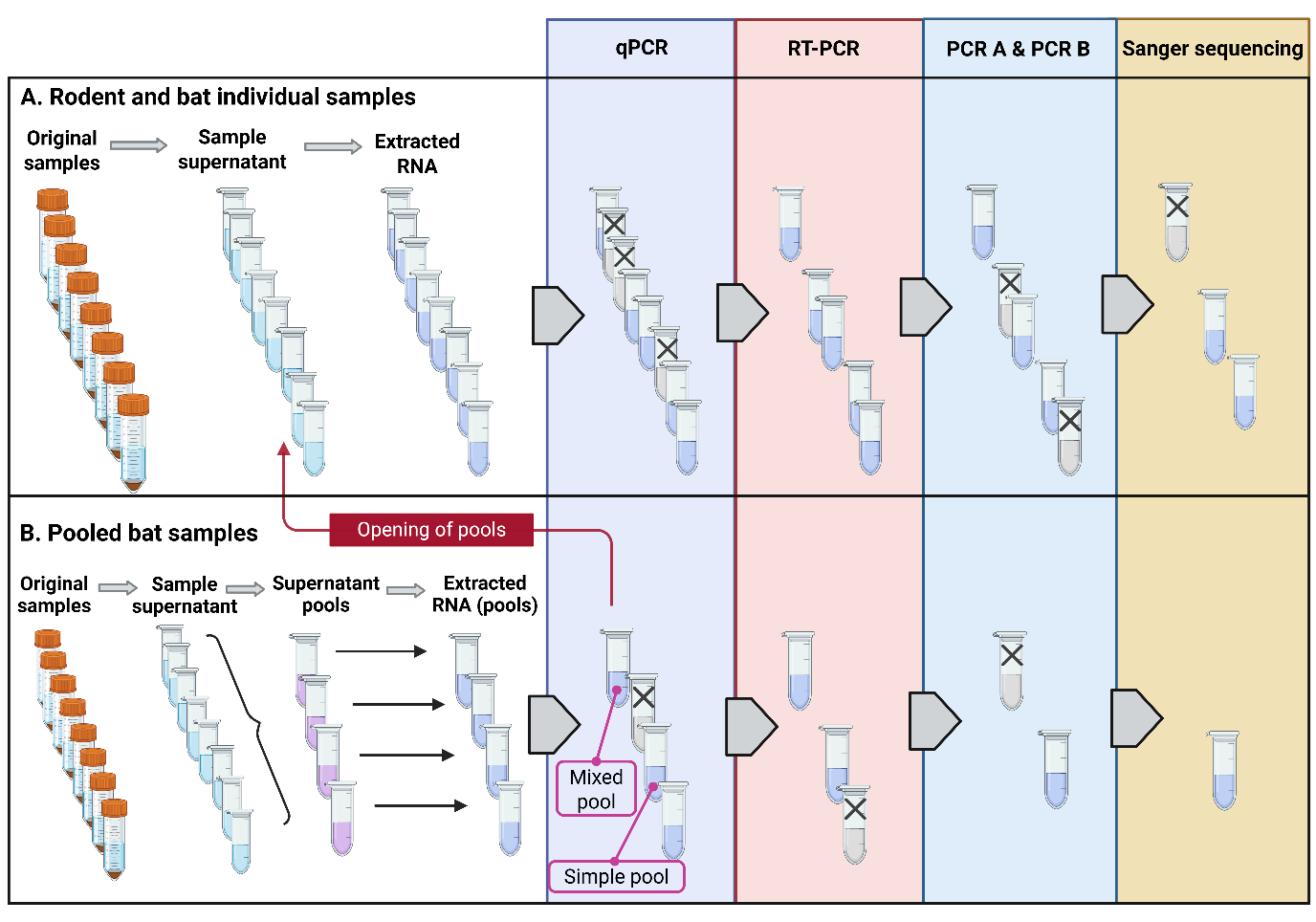
Terrestrial small mammals and part of bat samples were processed as individual samples (A), where samples were homogenised, and the cleared supernatant was used for RNA extraction. Due to a high number of Belgian bat guano, these samples were pooled for easier sample processing (B). Post screening qPCR, if the positive pool had individual samples from same sampling site collected during the same **sampling session** (simple pool), then the sample was carried forward as is, else (mixed pool) the individual samples from the pool were assessed with the workflow A to identify the actual positive sample. Figure created with BioRender.

**S4 Figure: Tanglegram with pruned tree and complete CDS tree**

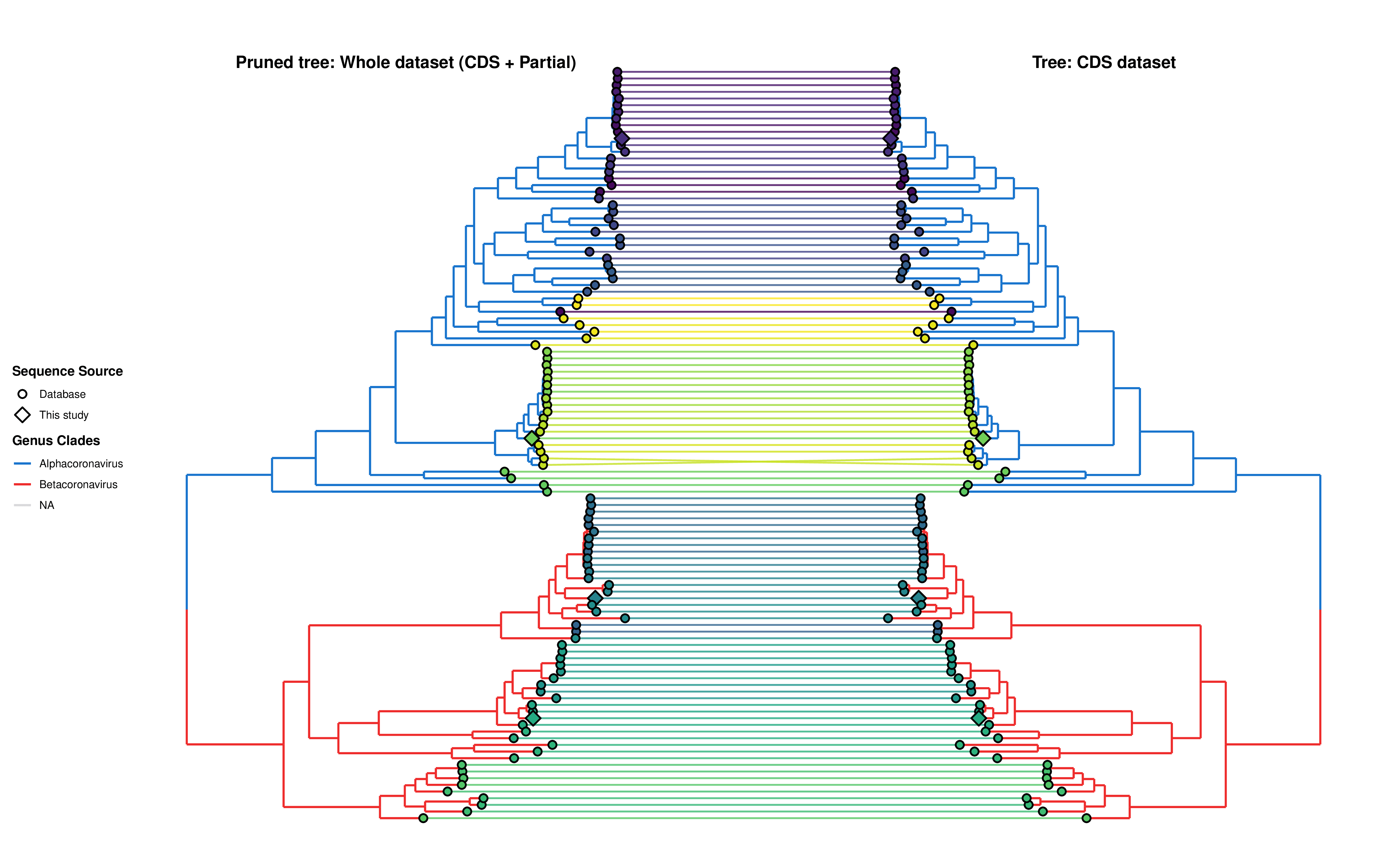

Assessment of topological differences were made using tanglegram, comparing the pruned tree and the tree made with partial RdRp sequences. More on the interpreted results can be found in Supplementary Material 4.
