## Supplementary material for "Detection and characterisation of alpha- and betacoronaviruses in rodents and bats from Germany, France, Belgium and Ireland"

**Supplementary Material 1: Protocol for amplification of the 476bp fragment of bat mitochondrial cytochrome B for identification**

A short region of cytochrome b covering 476bp was amplified and sequenced instead of the traditional 1140bp fragment due to possibility of degraded RNA in faeces.

The amplification reaction was prepared with GoTaq PCR mix, 2.5 mM of MgCl_2_, 200 nM of each primer and 5U of GoTaq and 1.5 µl of template. The conditions were as follows: denaturation at 94 °C for 5 minutes, 40 cycles with denaturation at 94 °C for 30 seconds, annealing at 50 °C for 30 seconds and extension at 72 °C for one minute, a final extension step of 72 °C for 10 minutes. Details can be found in S5 Table.

Post-amplification, the product was sequenced using the L14723_Cytb forward (Ducroz et al, 2001) and H15149_short reverse (Helm-Bychowski, et al. 1993) primers to identify the bats.

Ducroz JF, Volobouev V, Granjon L (2001) An Assessment of the systematics of arvicanthine rodents using mitochondrial DNA Sequences: evolutionary and biogeographical implications. J Mamm Evol 8: 173–206

Helm-Bychowski K, Cracraft J (1993) Recovering phylogenetic signal from DNA sequences: relationships within the corvine assemblage (class Aves) as inferred from complete sequences of the mitochondrial DNA cytochrome-b gene. Mol Biol Evol 10: 1196–1214

**Supplementary Material 2: Pre-processing of rodent colon tissue and bat faecal samples prior to viral RNA extraction**

**Rodent colon samples**

Sample preparation:

Colon samples stored in RNAlater at −20 °C were thawed prior to processing. Under a laminar-flow hood in a BSL-2 laboratory, approximately 5 mg of colon tissue free of faecal material was dissected and used for RNA extraction.

Tissue homogenisation and digestion:

Each 5 mg tissue sample was transferred into an Eppendorf tube containing sterile sand and two sterile glass beads together with 176 µl ACL buffer, 22 µl Proteinase K, and 210 µl phosphate-buffered saline (PBS). Mechanical homogenisation was performed using a TissueLyser II (QIAGEN, Germany) at 30 kHz for 2 min. After 1 min, the tube holders were inverted to ensure uniform disruption of the tissue. Following mechanical lysis, samples were incubated at 56 °C for 30 min to allow enzymatic digestion.

Lysate preparation:

Digested samples were centrifuged at 12,000 × *g* for 1 min to obtain a clarified lysate. A volume of 180 µl supernatant was used for viral RNA extraction.

**Bat faecal samples**

Removal of RNAlater:

Faecal samples stored in RNAlater at −20 °C were thawed to room temperature and centrifuged at 12,000 *× g* for 1 min. Excess RNAlater was removed manually by pipetting. Samples were washed once with 500 µl of 0.1 mM TE buffer and gently mixed. The suspension was incubated at room temperature for 1 h to dissolve residual RNAlater crystals. Following centrifugation at 12,000 × *g* for 1 min, the wash buffer was removed.

Faecal homogenisation:

After removal of the wash buffer, 400 µl EDTA solution was added to each sample. Samples were homogenised using a TissueLyser II (QIAGEN, Germany) at 30 kHz for 3 min. Homogenisation was paused midway through the run and the tubes were inverted to ensure uniform lysis. Samples were then centrifuged briefly at maximum speed for 1 min. Homogenates were either stored at −80 °C until processing of a complete extraction plate or immediately subjected to RNA extraction.

Pooling of Belgian bat samples:

Only Belgian bat samples were pooled prior to extraction. Individual samples were processed as described above before pooling. Pools were prepared by combining equal volumes of individual homogenates to produce a final volume of 200 µl.

Of the pools generated, 218 consisted of eight individual samples, six consisted of seven samples, and four consisted of nine samples (S3 Figure). Each sample was included in a single pool only. Individual homogenates were retained to enable follow-up testing of positive pools.

**Supplementary Material 3: Viral RNA extraction**

Viral RNA was extracted using the QIAamp 96 Virus QIAcube HT Kit and QIAcube HT extraction robot (QIAGEN, Germany) according to the manufacturer's instructions.

For rodent colon samples, 180 µl of clarified lysate was processed using the off-board lysis protocol. For bat faecal samples, 200 µl of homogenate was processed using the onboard lysis protocol, which included enzymatic lysis during extraction.

RNA was eluted in a final volume of 120 µl and used for downstream PCR screening, confirmatory assays, and sequencing analyses.

**Supplementary Material 4: Assessment of topological differences using Tanglegram**

Most study samples were represented by partial Sanger-derived RdRp sequences, whereas four samples (NCHA001531, BRDb538, PA20 and FASA10E) yielded complete RdRp coding sequences (CDS) through next-generation sequencing (NGS). To evaluate whether the inclusion of partial RdRp sequences influenced phylogenetic inference, a reference maximum-likelihood phylogeny was reconstructed using the four full-length study sequences together with reference full-length RdRp CDS retrieved from GenBank. A second phylogeny was reconstructed from a combined dataset containing both full-length RdRp sequences and partial RdRp fragments. Following tree inference, all partial sequences were removed by pruning, allowing direct comparison of taxa shared between the two phylogenies.

The alignment containing only full-length RdRp CDS sequences contained 5% non-informative sites, whereas the alignment including both full-length and partial RdRp sequences contained 21% non-informative sites. This increase in non-informative positions reduced the phylogenetic signal available for tree reconstruction.

Comparison of the resulting phylogenies using a tanglegram (S4 Figure) revealed minimal topological differences between the reference phylogeny inferred from full-length RdRp CDS sequences and the phylogeny inferred from the combined dataset. The tanglegram links corresponding taxa between the two trees, with crossing lines indicating differences in phylogenetic placement. Multiple taxa exhibited altered placements following inclusion of partial RdRp sequences, demonstrating that the addition of shorter fragments affected the inferred relationships among taxa.

These observations suggest that incorporation of partial RdRp fragments can influence phylogenetic reconstruction by increasing the proportion of non-informative sites within the alignment, thereby reducing phylogenetic resolution and altering tree topology.
